# Public Funding for AI in Canada 2011-2022: An equity-focused environmental scan

**DOI:** 10.1101/2025.01.31.635792

**Authors:** G. Attema, M. Mertz, A. Anawati, A. Austin, J. Bertrand, R. Jewett, E. Cameron

## Abstract

**Background:** In this new equity-driven landscape, if there is to be system-wide transformation in research funding allocation, indicators of funding allocations need to be explored. This environmental scan aims to understand how research funding for artificial intelligence (AI) has been allocated and distributed in Canada from 2011-2022. Using geographical representations, we describe and map research funding for Canadian researchers by publicly funded granting agencies and provide analyses for AI research spending since 2011

**Methodology:** We developed a rapid environmental scan to create a database of all AI funded projects from the following agencies: CIHR, NSERC, SSHRC, CRC, AMS, NFRF and CFI. Using publicly available research funding reporting and agency websites, we identified through title, keyword and project summary screening, AI projects in English and French for the years 2011-2022.

**Principal Findings:** A total of 4112 projects were identified, with the following information for each project recorded: title, year, institution, city, province, total funding, language, funding agency, funding program and primary investigator. A total of $384,933,265.74 million was allocated for publicly funded AI related projects in Canada from 2011-2022. Average funding per project was $93,612.18. The top three provinces with the most funding for all years are Ontario, Quebec, and British Columbia. The top three funding agencies by total amount for all years were NSERC at $155,267,817, CIHR at $136,594,644, and CFI at $58,317,627

**Conclusion:** This information can assist in accountability and understanding of Canada’s publicly funded research allocations, and provide information related to the distribution of such funds, thus informing equity policy strategies.

## Introduction

Artificial Intelligence research has been receiving increased attention over the last decade, as advances in ChatGPT, OpenAI and others have dramatically changed how and where we access information. While the potential for AI cannot be denied, with potential breakthroughs in information technology, health, commerce, and law to name a few, without equity and access, there lies the real potential to cleave open further the digital divide (1). Additionally, it has been shown that AI tools, whether used in healthcare or other sectors, can worsen existing biases, particularly those related to race, sex, and gender (2-6). AI in healthcare can improve accessibility and provide targeted interventions for underserved and vulnerable populations; however, biased datasets may perpetuate existing inequities (3,5). Mitigating biases, and addressing inequities requires a whole nation approach, particularly when it comes to research in this area. As such, our work began with the deceptively simple question, what research in AI in Canada has been funded?

## Public funding for research in Canada

Canadian researchers, particularly those who work in publicly funded institutions, rely primarily on federal funding allocated through the Tri-Agencies and other federally created programs (7). This research funding is comprised of three primary agencies: the Natural Sciences and Engineering Research Council of Canada (NSERC), the Canadian Institutes of Health Research (CIHR) and the Social Sciences and Humanities Research Council of Canada (SSHRC). While affiliated with the Tri-Agencies, Canada also has two additional funding programs that lie within and adjacent to the tri-agencies, the Canadian Foundation for Innovation (CFI), and the Canada Research Chairs program (CRC). Funding for these agencies comes from Canadian taxpayers, controlled by the Federal budget, and allocated generally through funding applications adjudicated using peer-review methods. While funding scoring differs between agencies, applications are scored using a merit-based system that looks at the following areas: quality of the application, quality of the research question, the researcher or research team, the feasibility and relevancy to Canadians, and the research environment (8). Additionally, federal research funding agencies have constructed initiatives to create a more equitable funding system in Canada, recognizing that systemic barriers can limit the potential for many skilled and excellent researchers (9,10). The Tri-Agency has committed to system-wide transformation in this area, launching ambitious targets for fair and equitable access to funding through their [Tri-Agency EDI Action Plan for 2018-2025](https://www.nserc-crsng.gc.ca/InterAgency-Interorganismes/EDI-EDI/Action-Plan_Plan-dAction_eng.asp?wbdisable=true), creating [new methods of peer review](https://cihr-irsc.gc.ca/e/52544.html) and committing to the [Dimensions Charter](https://www.nserc-crsng.gc.ca/InterAgency-Interorganismes/EDI-EDI/Dimensions-Charter_Dimensions-Charte_eng.asp).

## Equity driven landscape

In this new equity-driven landscape, if there is to be system-wide transformation in research funding allocation, indicators of funding allocations need to be explored. Our project is the first steps of this exploration by embarking on a robust environmental scan of research funding in Canada.

Given the recent explosion of AI funding, understanding where and what public funding has been allocated in this field helps to identify equity strategies in near-real time, and to establish important benchmarks for funding allocations moving forward. Using the environmental scan framework developed by Nagi et al. (11), we systematically searched funding databases in Canada, identifying projects which were related to AI. Funding databases and funding websites were searched beginning in June of 2023, and all funding related to AI for the period of 2011-2022 was collected. This information was then checked for accuracy and completeness and inputted into a database. Funding was collected based on fiscal year, with complete funding agencies reporting up to the end of fiscal year 2021-2022. Funding for the fiscal year 2022-2023 was available for the following agencies: AMS, CFI, CRC (all agencies) and CIHR. Funding from NSERC and SSHRC for the fiscal year 2022-2023 was not yet available. Results from this scan give us a first look at the landscape of AI funding in Canada from 2011-2022.

## Environmental scan methods

We began by defining Artificial Intelligence and all related terminology that would be used for inclusion. If a project matched to one or more of these AI terms it would be included in the dataset. After that we identified all research granting institutions that we were going to search for. Our focus was on government funding in publicly available granting databases from the period of 2011-2022. We did not search for industry related funding, or funding for people and projects outside of Canada. The period of 2011-2022 was chosen so that we could more effectively look at funding trends over time. Where available we used information found in the [Open Government Portal](https://open.canada.ca/en) and augmented it with searches on individual government and funding sites.

## Study inclusion criteria

Funded projects were included if they met the following Inclusion criteria: 1) Explicitly referred to AI and/or one of the following: machine learning, deep learning, natural language processing, computer vision, neural networks, expert (AI) systems, decision support systems, cognitive computing, robotics, intelligent (AI) agents, data mining, computer assisted diagnosis, computer assisted treatment, forecasting, telehealth, wearable technology, AI assisted precision medicine and/or health informatics and/or clinical decision support and/or electronic medical records, in the project title, keywords or project summary; 2) Funded between the years 2011-2022; 3) From the following funding agencies: CIHR, NSERC, SSHRC, Tri-Council (NFRF), AMS, CFI, and CRC; 4) Project takes place in Canada; 5) In English or in French; and 6) Grants and awards for individuals, teams, research chairs, interdisciplinary projects, graduates, postgraduates, and fellowships. Tri-Council (NFRF) are projects funded under the New Frontiers of Research Fund and are separated out because they are an interdisciplinary and distinct funding program. When initial project results from SSHRC were pulled, NFRF projects were also included as part of them, as the NFRF program is housed in the SSHRC database. We have moved all NFRF projects out of the SSHRC database and renamed them Tri-Council (NFRF).

## Data extraction strategy

Using publicly available funding reporting, we pulled all projects that were funded from the following agencies (AMS, CFI, CRC, SSHRC, NSERC, CIHR, Tri-Council (NFRF)) for the fiscal years 2011 to 2022. We then did a keyword search (AI and related terms) of all grants and kept those grants that matched. The following pieces of information from each project were recorded (Primary Investigator (PI), title, year awarded, agency, program, total amount awarded, language, institution, city, province). Once the data verification was completed, the remaining projects were downloaded to Excel where they were checked for quality and accuracy. Because of differences in reporting by funding agencies, some projects did not have a full set of information, and for those we went back to the funding websites and databases and performed additional searches to obtain this information. See Figure I for an overview of our search strategy and related results.

**Figure I:**
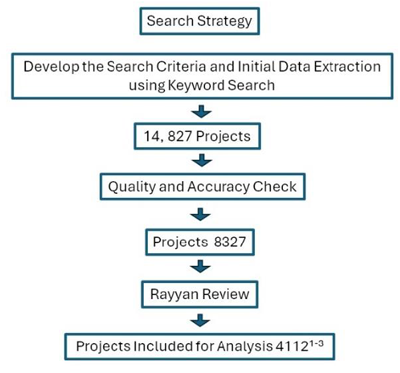
Search Strategy*, **, ***. 1) SSHRC data also included Tri-Council (NFRF) projects prior to the Rayyan review 2) CIHR numbers after the keyword search were unusually high, as CIHR projects in the Open Government Portal include far more text than any other agency. Because of this, many projects were included which upon review were not related to AI. 3) 158 projects from 2022-2023 were removed from the dataset. *Initial project search results by funding agency: NSERC 8058, CIHR 5037, SSHRC 950, CFI 569, AMS 58 **Project search result uploaded to Rayyan by funding agency: NSERC 3366, CIHR 3891, SSHRC 508, CFI 389, CRC 116, AMS 57 *CRC projects were removed from each tri-agency and placed in a standalone category ***Final included projects by funding agency: NSERC 3118, CIHR 528, SSHRC 199, CFI 170, Tri-Council (NFRF) 70, AMS 27 *CRC projects were added back to each tri-agency prior to analysis

## Data synthesis and presentation

Initial data pulled from tri-agency sites and funding websites yielded 14, 827 projects. After removal of duplicates and data clean up, 8342 projects remained. These projects were uploaded to [Rayyan](https://www.rayyan.ai/), where each one was reviewed by two members of the research team. A third reviewer was brought in to resolve any discrepancies. After the blinded review, 4112 projects were included in the final dataset, including 158 projects from the 2022-2023 fiscal year (which were removed prior to analysis). The total number of AI related projects for 2011-2022 per funding agency were as follows: AMS 27; CFI 170; SSHRC 199; CIHR 528; Tri-Council (NFRF) 70; NSERC 3118. CRC projects were added to their respective tri-council agencies (CIHR, NSERC and SSHRC).

## Results

Of the 4112 projects included in the dataset, total funding for all years is $384,933,265.74 million. The average funding for all years and all projects is $93,612.18. The largest number of projects funded comes from NSERC with 3118 projects totaling $155,267,817. One project was excluded as the total value of it far exceeded all other projects. *CGEn - Canada’s national facility for genome sequencing and analysis*, Naveed Aziz, 2022, Hospital for Sick Children, Toronto, Ontario, $48,900,667.00, CFI-2023 Major Science Initiatives Fund competition Projects were categorized as 1) Student/Post-doc/Training, 2) Individual Investigator (including new investigator), 3) Team, 4) Research Chair, 5) Interdisciplinary, 6) Institution, and 7) Other. Of the 4112 identified projects, 700 were Student/Post-doc/Training, 1985 were Individual Investigator, 267 were Team, 79 were Research Chairs, 243 were Interdisciplinary, 963 were Institute and 36 were Other. As far as the largest funding programs, NSERC’s Discovery Grants (Individual) had the most projects at 1674 projects, Engage Grants (University) had 673 projects, and The John F. Leaders Fund had 170 projects. The largest funder for graduate funding in AI was the Alexander G. Bell Doctoral Fund with 134 identified projects.

### Funding trends 2011-2022

Funding for AI related projects increased from $3,717,147 in 2011 to its highest recorded number in 2020 at $74,887,890 and falling from that high to $61,019,624 million in 2021. The total number of projects funded followed a similar trend, steadily increasing from 114 in 2011 to its peak in 2020 at 703, and then dipping to 577 in 2021. See Figure II

**Figure II:**
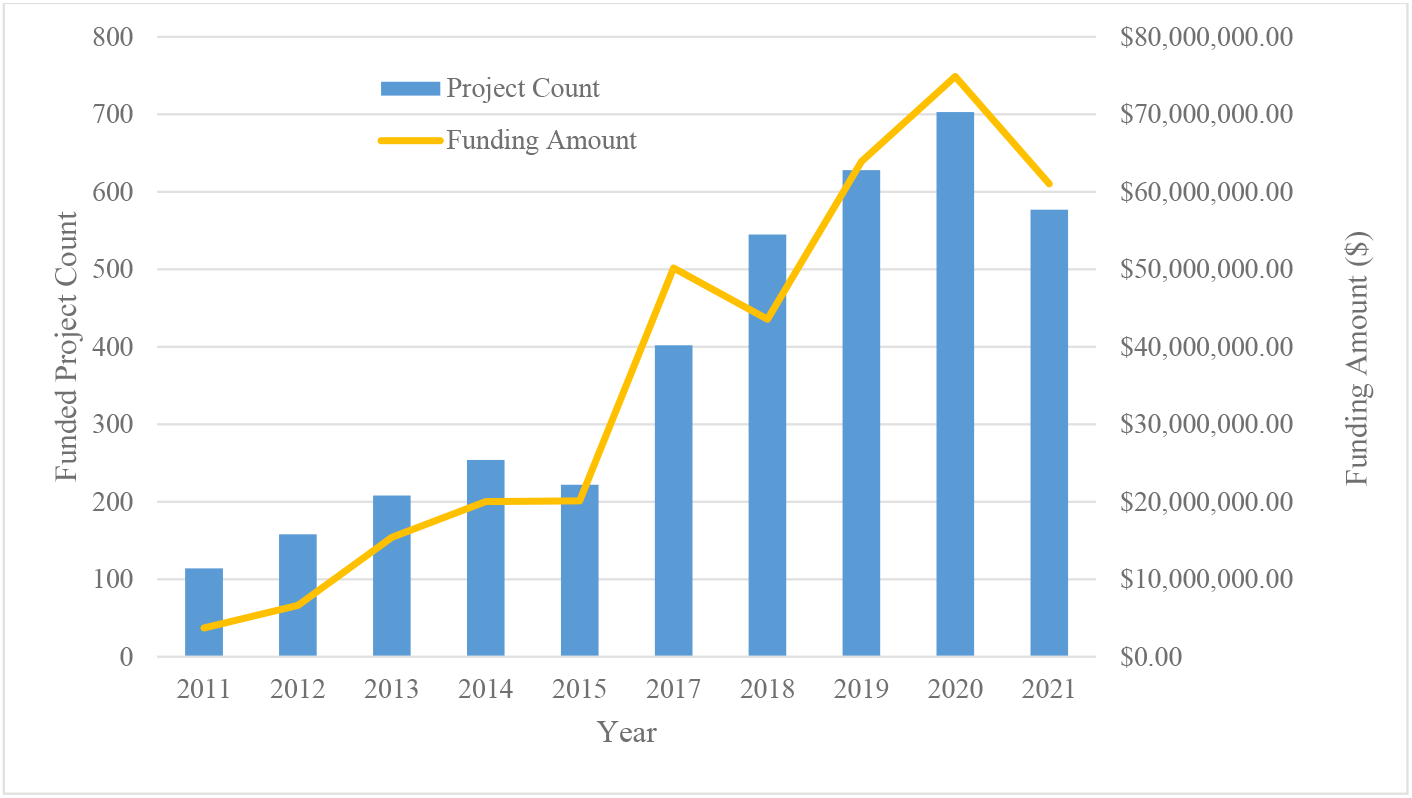
Project and funding overview 2011-2021.

### Canada Research Chairs in AI

We identified a total of 64 Canada Research Chairs (CRC) in AI from 2011-2022 (Table I). Total dollars allocated for CRC in AI was $47,900,000.00 million. There were 7(11%) CRCs in French, all located in Québec, at the Université de Montréal (6) and Université Laval (1), with 57 CRCs in English distributed across Canadian universities. A steady increase in CRCs in AI occurred between the years 2012-2016, with dramatic increases beginning in 2017-2022 with 54 CRCs allocated during that five-year span. Institutional allocations for CRCs are primarily concentrated in the largest universities in Canada including, the University of Toronto (10), the Université de Montréal (7), York (6), and McGill (6). Provincial distribution of CRCs shows the highest allocation of CRCs in Ontario (35) followed by Québec (18), and British Columbia (6).

**Table I:**
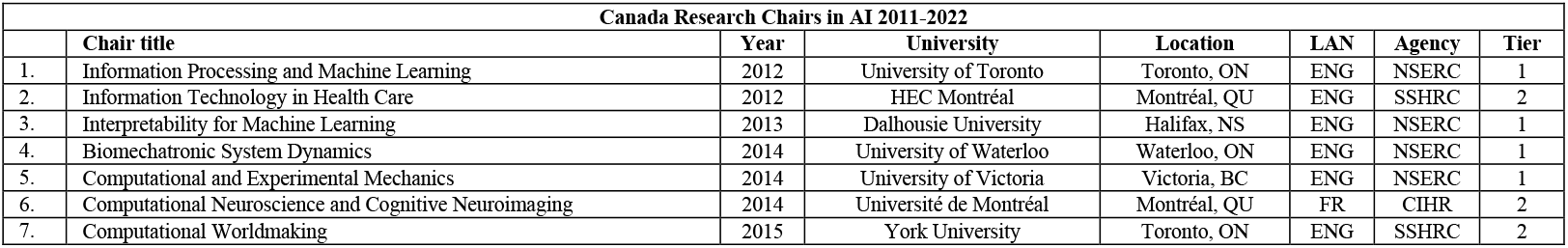
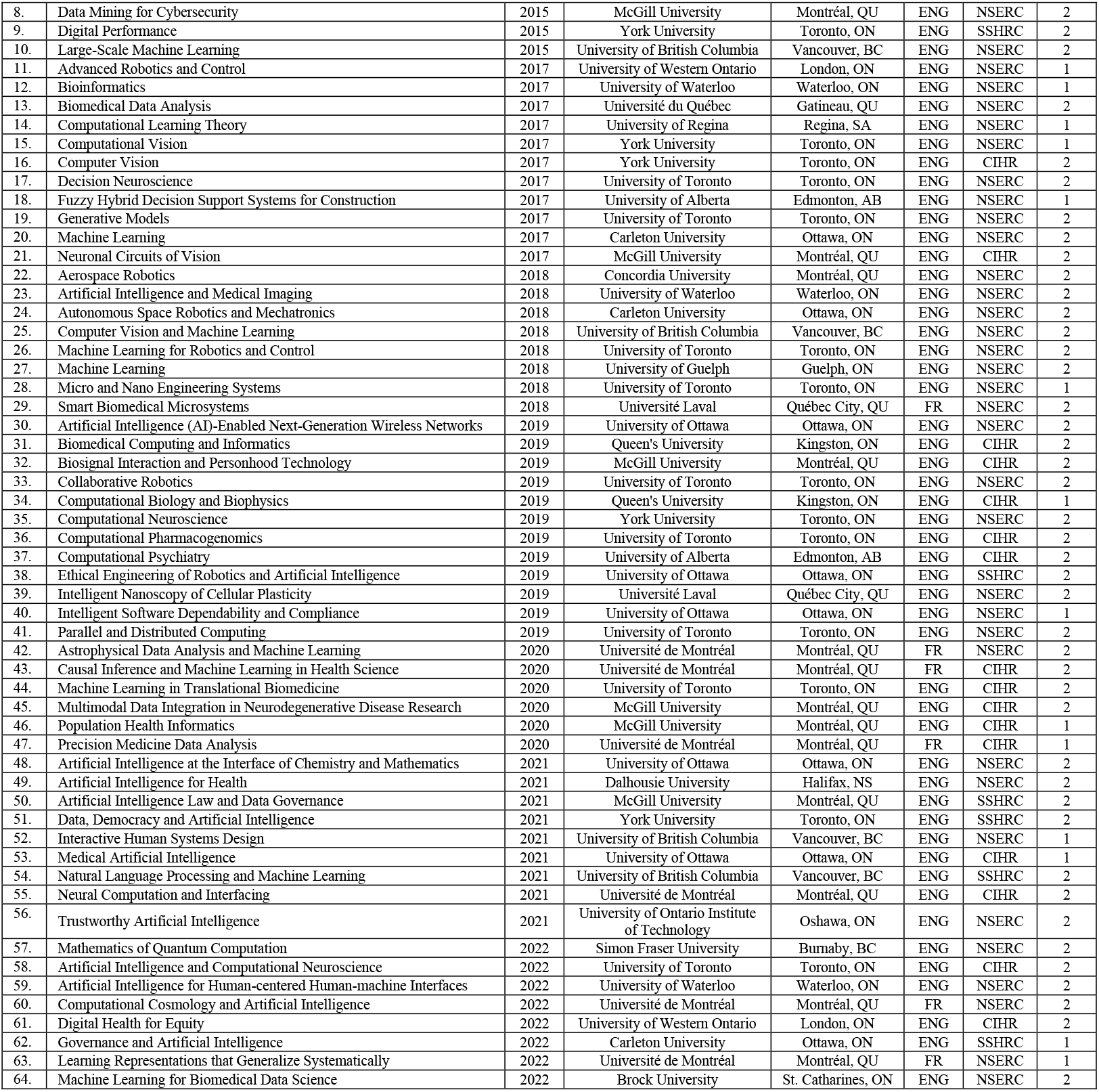
Canada Research Chairs in AI 2011-2022.

| Canada Research Chairs in AI 2011-2022 |  |  |  |  |  |  |  |
| --- | --- | --- | --- | --- | --- | --- | --- |
|  | Chair title | Year | University | Location | LAN | Agency | Tier |
| 1. | Information Processing and Machine Learning | 2012 | University of Toronto | Toronto, ON | ENG | NSERC | 1 |
| 2. | Information Technology in Health Care | 2012 | HEC Montréal | Montréal, QU | ENG | SSHRC | 2 |
| 3. | Interpretability for Machine Learning | 2013 | Dalhousie University | Halifax, NS | ENG | NSERC | 1 |
| 4. | Biomechatronic System Dynamics | 2014 | University of Waterloo | Waterloo, ON | ENG | NSERC | 1 |
| 5. | Computational and Experimental Mechanics | 2014 | University of Victoria | Victoria, BC | ENG | NSERC | 1 |
| 6. | Computational Neuroscience and Cognitive Neuroimaging | 2014 | Université de Montréal | Montréal, QU | FR | CIHR | 2 |
| 7. | Computational Worldmaking | 2015 | York University | Toronto, ON | ENG | SSHRC | 2 |
| 8. | Data Mining for Cybersecurity | 2015 | McGill University | Montréal, QU | ENG | NSERC | 2 |
| 9. | Digital Performance | 2015 | York University | Toronto, ON | ENG | SSHRC | 2 |
| 10. | Large-Scale Machine Learning | 2015 | University of British Columbia | Vancouver, BC | ENG | NSERC | 2 |
| 11. | Advanced Robotics and Control | 2017 | University of Western Ontario | London, ON | ENG | NSERC | 1 |
| 12. | Bioinformatics | 2017 | University of Waterloo | Waterloo, ON | ENG | NSERC | 1 |
| 13. | Biomedical Data Analysis | 2017 | Université du Québec | Gatineau, QU | ENG | NSERC | 2 |
| 14. | Computational Learning Theory | 2017 | University of Regina | Regina, SA | ENG | NSERC | 1 |
| 15. | Computational Vision | 2017 | York University | Toronto, ON | ENG | NSERC | 1 |
| 16. | Computer Vision | 2017 | York University | Toronto, ON | ENG | CIHR | 2 |
| 17. | Decision Neuroscience | 2017 | University of Toronto | Toronto, ON | ENG | NSERC | 2 |
| 18. | Fuzzy Hybrid Decision Support Systems for Construction | 2017 | University of Alberta | Edmonton, AB | ENG | NSERC | 1 |
| 19. | Generative Models | 2017 | University of Toronto | Toronto, ON | ENG | NSERC | 2 |
| 20. | Machine Learning | 2017 | Carleton University | Ottawa, ON | ENG | NSERC | 2 |
| 21. | Neuronal Circuits of Vision | 2017 | McGill University | Montréal, QU | ENG | CIHR | 2 |
| 22. | Aerospace Robotics | 2018 | Concordia University | Montréal, QU | ENG | NSERC | 2 |
| 23. | Artificial Intelligence and Medical Imaging | 2018 | University of Waterloo | Waterloo, ON | ENG | NSERC | 2 |
| 24. | Autonomous Space Robotics and Mechatronics | 2018 | Carleton University | Ottawa, ON | ENG | NSERC | 2 |
| 25. | Computer Vision and Machine Learning | 2018 | University of British Columbia | Vancouver, BC | ENG | NSERC | 2 |
| 26. | Machine Learning for Robotics and Control | 2018 | University of Toronto | Toronto, ON | ENG | NSERC | 2 |
| 27. | Machine Learning | 2018 | University of Guelph | Guelph, ON | ENG | NSERC | 2 |
| 28. | Micro and Nano Engineering Systems | 2018 | University of Toronto | Toronto, ON | ENG | NSERC | 1 |
| 29. | Smart Biomedical Microsystems | 2018 | Université Laval | Québec City, QU | FR | NSERC | 2 |
| 30. | Artificial Intelligence (AI)-Enabled Next-Generation Wireless Networks | 2019 | University of Ottawa | Ottawa, ON | ENG | NSERC | 2 |
| 31. | Biomedical Computing and Informatics | 2019 | Queen's University | Kingston, ON | ENG | CIHR | 2 |
| 32. | Biosignal Interaction and Personhood Technology | 2019 | McGill University | Montréal, QU | ENG | CIHR | 2 |
| 33. | Collaborative Robotics | 2019 | University of Toronto | Toronto, ON | ENG | NSERC | 2 |
| 34. | Computational Biology and Biophysics | 2019 | Queen's University | Kingston, ON | ENG | CIHR | 1 |
| 35. | Computational Neuroscience | 2019 | York University | Toronto, ON | ENG | NSERC | 2 |
| 36. | Computational Pharmacogenomics | 2019 | University of Toronto | Toronto, ON | ENG | CIHR | 2 |
| 37. | Computational Psychiatry | 2019 | University of Alberta | Edmonton, AB | ENG | CIHR | 2 |
| 38. | Ethical Engineering of Robotics and Artificial Intelligence | 2019 | University of Ottawa | Ottawa, ON | ENG | SSHRC | 2 |
| 39. | Intelligent Nanoscopy of Cellular Plasticity | 2019 | Université Laval | Québec City, QU | ENG | NSERC | 2 |
| 40. | Intelligent Software Dependability and Compliance | 2019 | University of Ottawa | Ottawa, ON | ENG | NSERC | 1 |
| 41. | Parallel and Distributed Computing | 2019 | University of Toronto | Toronto, ON | ENG | NSERC | 2 |
| 42. | Astrophysical Data Analysis and Machine Learning | 2020 | Université de Montréal | Montréal, QU | FR | NSERC | 2 |
| 43. | Causal Inference and Machine Learning in Health Science | 2020 | Université de Montréal | Montréal, QU | FR | CIHR | 2 |
| 44. | Machine Learning in Translational Biomedicine | 2020 | University of Toronto | Toronto, ON | ENG | CIHR | 2 |
| 45. | Multimodal Data Integration in Neurodegenerative Disease Research | 2020 | McGill University | Montréal, QU | ENG | CIHR | 2 |
| 46. | Population Health Informatics | 2020 | McGill University | Montréal, QU | ENG | CIHR | 1 |
| 47. | Precision Medicine Data Analysis | 2020 | Université de Montréal | Montréal, QU | FR | CIHR | 1 |
| 48. | Artificial Intelligence at the Interface of Chemistry and Mathematics | 2021 | University of Ottawa | Ottawa, ON | ENG | NSERC | 2 |
| 49. | Artificial Intelligence for Health | 2021 | Dalhousie University | Halifax, NS | ENG | NSERC | 2 |
| 50. | Artificial Intelligence Law and Data Governance | 2021 | McGill University | Montréal, QU | ENG | SSHRC | 2 |
| 51. | Data, Democracy and Artificial Intelligence | 2021 | York University | Toronto, ON | ENG | SSHRC | 2 |
| 52. | Interactive Human Systems Design | 2021 | University of British Columbia | Vancouver, BC | ENG | NSERC | 1 |
| 53. | Medical Artificial Intelligence | 2021 | University of Ottawa | Ottawa, ON | ENG | CIHR | 1 |
| 54. | Natural Language Processing and Machine Learning | 2021 | University of British Columbia | Vancouver, BC | ENG | SSHRC | 2 |
| 55. | Neural Computation and Interfacing | 2021 | Université de Montréal | Montréal, QU | ENG | CIHR | 2 |
| 56. | Trustworthy Artificial Intelligence | 2021 | University of Ontario Institute of Technology | Oshawa, ON | ENG | NSERC | 2 |
| 57. | Mathematics of Quantum Computation | 2022 | Simon Fraser University | Burnaby, BC | ENG | NSERC | 2 |
| 58. | Artificial Intelligence and Computational Neuroscience | 2022 | University of Toronto | Toronto, ON | ENG | CIHR | 2 |
| 59. | Artificial Intelligence for Human-centered Human-machine Interfaces | 2022 | University of Waterloo | Waterloo, ON | ENG | NSERC | 2 |
| 60. | Computational Cosmology and Artificial Intelligence | 2022 | Université de Montréal | Montréal, QU | FR | NSERC | 2 |
| 61. | Digital Health for Equity | 2022 | University of Western Ontario | London, ON | ENG | CIHR | 2 |
| 62. | Governance and Artificial Intelligence | 2022 | Carleton University | Ottawa, ON | ENG | SSHRC | 1 |
| 63. | Learning Representations that Generalize Systematically | 2022 | Université de Montréal | Montréal, QU | FR | NSERC | 1 |
| 64. | Machine Learning for Biomedical Data Science | 2022 | Brock University | St. Catharines, ON | ENG | NSERC | 2 |

### Provinces most funded

Looking at total funding dollars by province reveals Ontario with the largest amount at just over 173 million dollars, Québec with just over 87 million dollars and British Columbia with just over 59 million dollars. Provinces with the least amount of total funding include Prince Edward Island with just over 450 thousand dollars, New Brunswick with 1.6 million dollars and Newfoundland and Labrador with 2.2 million dollars. Expressed as percentages of the total funding allocated, 82% of all funding was allocated to three provinces, Ontario (45%), Québec (22%), and British Columbia (15%). Total funding for Maritime provinces was less than 4%, with three provinces: Newfoundland and Labrador (0.52%), New Brunswick (0.42%), and Prince Edward Island (0.12%) each having less than 1%. Total amounts for each province for each year can be found in Table II. Funding allocations were geographically mapped by institute location (city) and can be found in Image I.

**Table II:**
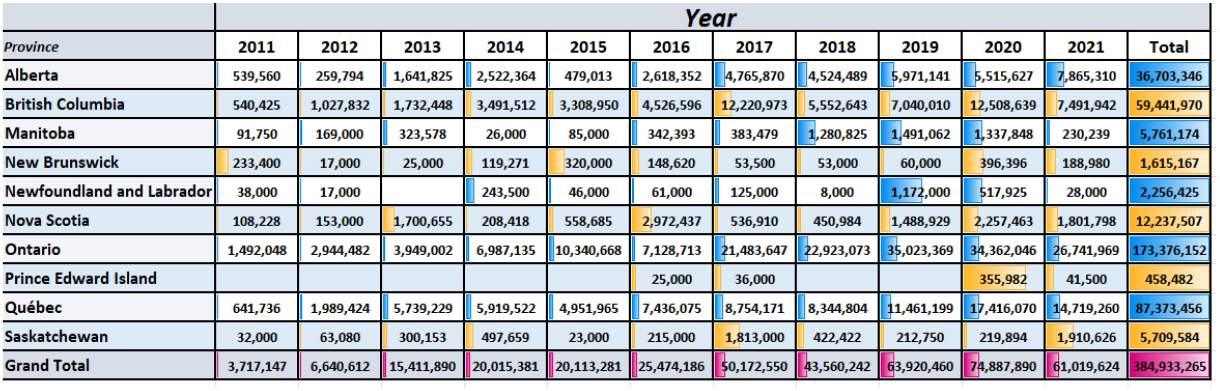
AI funding in Canada by Province: Total (2011-2021) and by year.

**Image I:**
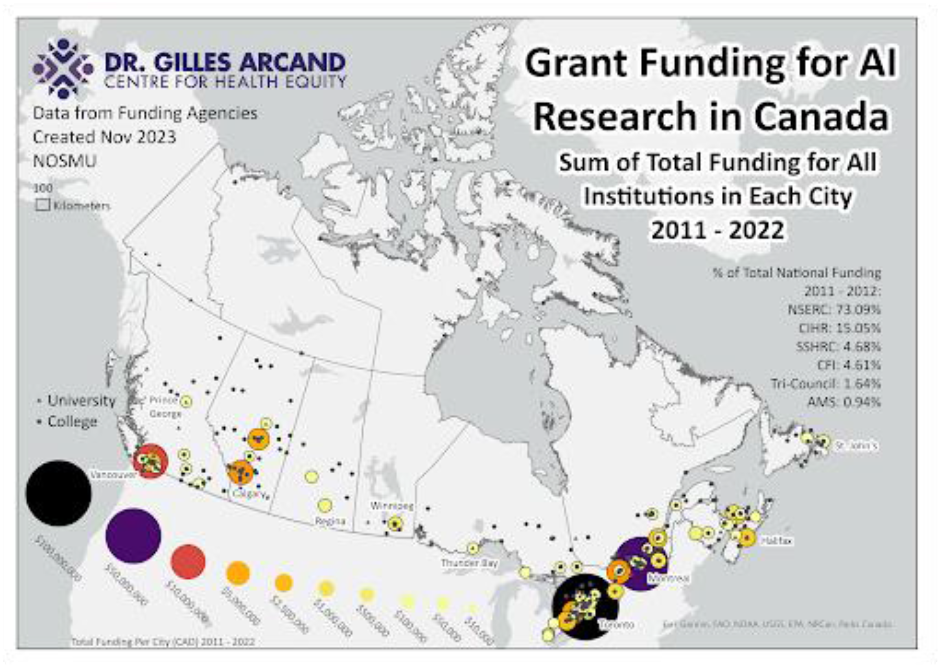
Geographical representation of funding allocation.

### Institutions most funded

A look at the top institutions who received AI funding unsurprisingly includes Canada’s largest universities, along with its major research centres, institutes, and hospitals. The University of Toronto received the most amount of total funding dollars at just over 38.6 million, followed by the University of British Columbia at nearly 37 million. McGill University received just over 27 million dollars in funding, and the University of Calgary and York University with 19.6 million and nearly 19 million respectively. See Table III

**Table III:** Top Funded Institutions.

| Rank | Institution | City | Province | Total Funding |
| --- | --- | --- | --- | --- |
| 1 | University of Toronto | Toronto | Ontario | \$38,630,808.67 |
| 2 | University of British Columbia | Vancouver | British Columbia | \$36,205,715.17 |
| 3 | McGill University | Montréal | Québec | \$27,549,243.32 |
| 4 | University of Calgary | Calgary | Alberta | \$19,647,147.78 |
| 5 | York University | Toronto | Ontario | \$18,936,584.00 |
| 6 | University of Waterloo | Waterloo | Ontario | \$17,727,854.66 |
| 7 | University of Alberta | Edmonton | Alberta | \$15,563,925.65 |
| 8 | University of Ottawa | Ottawa | Ontario | \$13,323,621.32 |
| 9 | Simon Fraser University | Burnaby | British Columbia | \$12,095,568.00 |
| 10 | Université de Montréal | Montréal | Québec | \$11,601,479.00 |
| 11 | Université Laval | Laval | Québec | \$10,720,200.97 |
| 12 | Dalhousie University | Halifax | Nova Scotia | \$10,369,976.00 |
| 13 | University of Victoria | Victoria | British Columbia | \$9,308,774.66 |
| 14 | Queen's University | Kingston | Ontario | \$8,804,387.63 |
| 15 | University Health Network | Toronto | Ontario | \$8,670,436.00 |
| 16 | Université de Sherbrooke | Sherbrooke | Québec | \$7,874,550.66 |
| 17 | Unity Health Toronto | Toronto | Ontario | \$7,110,188.00 |
| 18 | University of Western Ontario | London | Ontario | \$6,862,378.00 |
| 19 | McMaster University | Hamilton | Ontario | \$6,434,992.08 |
| 20 | École Polytechnique de Montréal | Montréal | Québec | \$5,689,811.00 |
| 21 | Concordia University | Montréal | Québec | \$5,424,083.00 |
| 22 | University of Manitoba | Winnipeg | Manitoba | \$5,309,736.32 |
| 23 | Hospital for Sick Children | Toronto | Ontario | \$4,393,692.00 |
| 24 | Sheridan Institute of Technology and Advanced Learning | Oakville | Ontario | \$4,120,487.00 |
| 25 | Carleton University | Ottawa | Ontario | \$4,076,237.00 |
| 26 | Research Institute of the McGill University Health Centre | Montréal | Québec | \$3,968,106.00 |
| 27 | University of Regina | Regina | Saskatchewan | \$3,687,465.00 |
| 28 | Toronto Metropolitan University | Toronto | Ontario | \$3,237,706.00 |
| 29 | University of Guelph | Guelph | Ontario | \$3,161,535.00 |
| 30 | Memorial University of Newfoundland | St. John's | Newfoundland | \$2,256,425.00 |

### Funding by language

An examination of AI funding by language, reveals a paucity of funding for French language projects. We identified only 29 French projects for all years, which represents 0.7% of the total number of projects. The total value of French projects was $6,602,976, or 1.71% of the total amount allocated for AI funding. The breakdown of this funding by agency total funding dollars shows CIHR as the top funder with 8 projects worth just over 3.4 million, followed closely by NSERC with 10 projects valued at just over 1.2 million. The Canada Research Chairs program had the highest French language representation, with 7 (11%) French Research Chairs of a total of 64. See Box I.

#### Box I

**French language funded projects**

Box I: French language funded projects
| French Language Funded Projects in AI 2011-2022 |  |
| --- | --- |
| Total number of projects (% of total funding) | 29 (.71%) |
| Total value of projects (% of total funding) | \$6,602,976.00 (1.72%) |
| Breakdown of projects by funding agency (total amount for all years) | SSHRC-3 (\$138,652.00)<br>CIHR-9 (\$3,473,488.00)<br>CFI-8 (\$1,753,484.00)<br>NSERC-12 (\$1,237,352.00) |

## Discussion

Funding and total number of funded projects for AI research in Canada among these funding sources has steadily increased since 2011, peaking in 2020. The total value of funding dollars in AI research has dropped, from its peak of 74 million and 703 projects in 2020, to 61 million and 577 projects in 2021. This 8% drop might signal the tapering of funding for AI related research or could be akin to AI related funding in 2018 which saw an overall decrease of just over 6.5 million from the previous year. The largest increase in year-over-year funding was in 2017, where AI funding nearly doubled from just over 25 million in 2016 to just over 50 million in 2017. Other factors, such as overall federal funding for research, effects of the COVID-19 pandemic on AI research, and other system-wide influences, which are beyond the scope of this paper, need to be considered when interpreting these figures and could be useful areas of further research. The concentration of AI funding to large urban cities and institutes is not surprising, but given the government’s commitments to equity, diversity and inclusion in research funding, rationale for allocations and other equity focused research is needed. There appears to be a movement towards equity in French language funding, but only in one program, the Canada Research Chairs, where 11% of chair ships in AI were awarded to French language chairs. In all other measures, French language projects were underrepresented and totaled less than 2% of all funding allocated. As AI will factor largely in the lives of all Canadians, deeper analysis of the scope and type of funded AI research in Canada is needed, with this environmental scan representing the first step in this important work.

### Limitations and future areas of research

There are a few limitations with this scan. First, we were only able to identify publicly available data, and because of that we could not get a complete dataset for the fiscal year 2022-2023. We also relied on the quality of both the inputting of the data, but also the robustness of the dataset programs themselves. This required us to manually input many projects, a stark reminder that the quality of automation is only as good as the data that is used to create it. Given the delay in reporting from some agencies, we cannot extend any trend analysis beyond the fiscal year of 2021-22 and given the speed at which AI has captivated the world, we stipulate that there is important data that cannot be included. We also limited our data to publicly available research funding and did not include industry, commercialization and innovation data.

Another limitation is the amount of information we were able to pull from each project. Some databases allow for keywords and project abstracts as part of the information included, while others only listed project titles. Where information was limited, we tried to pull out other available information, but it was not possible or practical to do this for all funded projects in all agencies.

Future areas of research can continue to examine research funding allocations in Canada, with the goal to align funding allocations with equity, diversity and inclusion goals set out by the federal funding agencies. Another area of future research could be more detailed examinations of the geographical distribution of research funding across Canada.

## Conclusion

In Canada, research in the post-secondary setting is largely funded by the federal government, with the goal of benefiting all Canadians. The Tri-Agency funding agencies have publicly identified inequalities and a lack of diversity and inclusion in both who become researchers and how they are funded (10). As part of a national effort to address this, the tri-agencies have committed to system-wide reviews and reforms (9, 10). We argue that central to this system-wide transformation is the knowledge of who receives funding, for what purpose and in what setting. Given the importance, relevancy and need for research in Artificial Intelligence, this environmental scan of funded projects is a first step in understanding research funding allocation so that we may collectively work towards equity-driven solutions. It is our hope that others work alongside us, and use this approach for other research fields, and in doing so provide analysis and transparency, two key aspects required for true system-wide transformation.

## List of Abbreviations

AMS: Associated Medical Services, a funding agency in Canada which focuses on research in compassion and technology-driven care and the history of medicine. https://www.ams-inc.on.ca/CFI-Canadian Foundation for Innovation, a non-profit corporation that invests in research infrastructure at Canadian universities, colleges, research hospitals and non-profit research institutes. https://www.innovation.ca/
CIHR: Canadian Institutes of Health Research, Canada’s federal funding agency for health research, comprised of 13 institutes (Aging; Cancer Research; Circulatory and Respiratory Health; Gender and Health; Genetics; Health Services and Policy Research; Human Development, Child and Youth Health; Indigenous Peoples’ Health; Infection and Immunity; Musculoskeletal Health and Arthritis; Neurosciences, Mental Health and Addiction; Nutrition, Metabolism and Diabetes; Population and Public Health). Part of the federal tri-agency granting councils of Canada. https://cihr-irsc.gc.ca
CRC: Canada Research Chairs, part of the federal tri-agency granting councils of Canada and is part of a national strategy to attract and retain top research talent in Canada through two streams of Research Chairs, tier 1 and tier 2. Research chairs are allocated based on the level of tri-agency funding for institutions, with Tier 1 Chairs valued at $1,400, 000 and Tier 2 Chairs valued at $500,000. https://www.chairs-chaires.gc.ca/home-accueil-eng.aspx
NFRF: New Frontiers in Research Fund, is part of the federal tri-agency granting councils of Canada, supporting world-leading, interdisciplinary research that is international, transformative and rapid-response. They have three streams to support their vision, Exploration; International; Transformation, and can create rapid-response special calls. https://www.sshrc-crsh.gc.ca/funding-financement/nfrf-fnfr/index-eng.aspx
NSERC: National Sciences and Engineering Research Council of Canada, is part of the federal tri-agency granting councils of Canada and supports research to advance scientific and technical breakthroughs. Their research priority areas include, Aerospace; Automotive; Environmental Science and Agriculture; Forestry and Wood Products Research; Life Sciences and Related Technologies; Information and Communications Technologies; Natural Resources and Energy; Northern Research; Manufacturing; Oil Sands and Heavy Oil; and Water-Related Research. https://www.nserc-crsng.gc.ca/index_eng.asp
SSHRC: Social Sciences and Humanities Research Council of Canada, is part of the federal tri-agency granting council of Canada and supports research and training in the humanities and social sciences. Their mandate is to develop the talent of Canadian researchers in building knowledge to better understand society and driving innovations. https://www.sshrc-crsh.gc.ca/home-accueil-eng.aspx

## Acknowledgements

We would like to thank the members of the AI-North community for their support of this work.

## Notes

https://open.canada.ca/en

### Competing Interest Statement

The authors have declared no competing interest.

## References

1. van Dijk, J. (2020). The digital divide. John Wiley & Sons, Cambridge

2. Char, D.S., Abramoff, M.D., & Feudtner, C. (2020). Identifying ethical considerations for machine learning healthcare applications. Am J Bioeth 20:7–17. doi:10.1080/15265161.2020.1819469

3. Fisher, S., & Rosella, L.C. (2022). Priorities for successful use of artificial intelligence by public health organizations: a literature review. BMC Public Health 22:2146. 10.1186/s12889-022-14422-z

4. He, J., Baxter, S.L., Xu, J., Xu. J.I., Zhou, X., & Zhang, K. (2019). The practical implementation of artificial intelligence technologies in medicine. Nat Med 25:30–36. 10.1038/s41591-018-0307-0

5. Rajpurkar, P., Chen, E., Banerjee, O., & Topol, E.J. (2022). AI in health and medicine. Nat Med 28:31–38. 10.1038/s41591-021-01614-0

6. Weins, J., Saria, S., Sendak, M., Ghassemi, M., Liu, V.X., Doshi-Velez, F., Jung, K., Heller, K., Kale, D., Saeed, M., Ossorio, P.N., Thadaney-Israni, S., & Goldenberg, A. (2019). Do no harm: a roadmap for responsible machine learning for health care. Nat Med 25:1337–1340. 10.1038/s41591-019-0548-6

7. Government of Canada. (nd). Open government. https://open.canada.ca/en Accessed June to October 2023

8. Government of Canada. (2022). Tri-Agency interdisciplinary peer review committee. https://cihr-irsc.gc.ca/e/52544.html Accessed 05 December 2023

9. Government of Canada. (2018). Tri-Agency equity, diversity and inclusion action plan for 2018-2025. https://www.nserc-crsng.gc.ca/_doc/EDI/EDI-ActionPlan-EN.pdf Accessed 05 December 2023

10. Government of Canada. (2019). Dimensions: equity, diversity and inclusion Canada. http://www.nserc-crsng.gc.ca/InterAgency-Interorganismes/EDI-EDI/Dimensions-Charter_Dimensions-Charte_eng.asp Accessed 05 December 2023

11. Nagi, R., Rogers Van Katwyk, S., & Hoffman, S.J. (2020). Using a rapid environmental scan methodology to map country-level global health research expertise in Canada. Health Research Policy and Systems 18:37 10.1186/s12961-020-0543-x

